# RNA sequencing identifies lung cancer lineage and facilitates drug repositioning

**DOI:** 10.1101/2023.01.18.524544

**Authors:** Longjin Zeng, Longyao Zhang, Lingchen Li, Xingyun Liao, Chenrui Yin, Lincheng Zhang, Xiewan Chen, Jianguo Sun

## Abstract

Breakthrough therapies recently improve survival in lung adenocarcinoma (LUAD), yet we still lack a paradigm to support prospective confirmation. To classify three clusters including bronchioid, neuroendocrine, and squamoid, the non-negative matrix factorization (NMF) algorithm was first performed at The Cancer Genome Atlas (TCGA), Clinical Proteomic Tumor Analysis Consortium (CPTAC), Gene Expression Omnibus (GEO) and preclinical models. In terms of prognostic value, the squamoid cluster is of poor prognostic factor. The neuroendocrine cluster is characterized by STK11 mutations and 14q13.3 amplifications. From an immunological perspective, the bronchioid cluster is considered an immune activation because of the highest immune-related genetic perturbation. Further analysis is the estimation of the relative cell abundance of the tumor microenvironment (TME), specific cell types can be reflected among three clusters. Meanwhile, the higher portion of immune cell infiltration belonged to bronchioid and squamous, not neuroendocrine cluster. Taken together, the neuroendocrine cluster show resistance to PD-L1 blockade. While pemetrexed or platinum-based therapies are suitable for bronchioid and squamoid clusters, respectively. Our emphasis was on phenotype-based action to explore compounds. Large-scale drug sensitivity databases including ConnectivityMap (CMAP), Cancer Cell Line Encyclopedia (CCLE), and Genomics of Drug Sensitivity in Cancer (GDSC) were analyzed. MEK inhibitors exhibited resistance in the bronchioid, while sensitive to the squamous cluster. Dinaciclib and alvocidib showed similar activity and sensitivity in the neuroendocrine cluster. A lineage factor named KLF5 recognized by two networks could be suppressed by verteporfin. This work adds to the knowledge of the lung cancer lineage and facilitates drug repositioning.

## 1. Introduction

It is widely generally accepted that the development of advanced non-small cell lung cancer (NSCLC) is based on genetic mutations, some of which are driver mutations. Tyrosine kinase inhibitor therapy is used to target mutated oncogenes.^1,2^ However, a large proportion of the population lacks a targetable driver and is referred to as “driver-negative”.^2^ In recent years, anti-programmed death 1 (PD-1/PD-L1) immune checkpoint inhibitor (ICI) therapies become the keystone of first-line therapy for driver-negative NSCLC.^2^ Preclinical evidence suggests that the addition of anti-PD-1/PD-L1 therapies improved prognosis in patients treated with combined pemetrexed and platinum and showed great potential value in nonsquamous NSCLC treatment.^2,3^ The TME suitable for ICI treatments is defined as “hot-immune” with higher cytotoxic T cell infiltration and tumor mutation burden (TMB).^4^

TCGA revealed the existence of three transcriptional subtypes in LUAD, including bronchioid/terminal respiratory unit (TRU), magnoid/proximal-proliferative (PP), and squamoid/proximal-inflammatory (PI).^5^ TCGA group proved TRU subtype was associated with a better prognosis and was enriched for EGFR mutations. The loss function of KRAS and STK11 occurred frequently in the PP subtype. While the PI subtype was described as a co-occurrence of TP53 and NF1. Importantly, PP and PI patients have a poorer prognosis compared to TRU, which may be due to their highly proliferative characteristics.^5,6^ Ringnér et al. reported a technical bias in the TCGA subtype and that metagenes may be a valid improvement.^6^ Moreover, we think it is counterintuitive because lung cancer lineage infidelity should be less proportional.^7^

Recent studies have shown that the transcriptome determines the fate of lung cancer lineage rather than the genome.^8^ In fact, lung cancer lineage has received attention in preclinical models.^9^ We suppose lung cancer lineage infidelity may acquire therapeutic resistance. In this study, we investigated the relationship between lung cancer lineage, immunity, and drug utilization.

## 2. Materials and Methods

### 2.1 Clinical cohorts and preprocessing

For the main analysis, a total of 604 patients with stage IB-IIIA LUAD in the TCGA-LUAD (n = 320), GSE72094 (n = 207), and CPTAC-LUAD (n = 77) cohorts were included. UCSC Xena website was used to download TCGA level three RNA-seq data (Illumina HiSeq 2000).^5,10,11,12^ RNA-seq count data were transformed into Transcripts Per Million (TPM) for analysis. The expression datasets were downloaded from the GEO database (**https://www.ncbi.nlm.nih.gov/geo/**) and quantile-normalized. Then transformed using log_(x+1)_ and log_2_ for TCGA and GEO respectively. Updated clinical information, copy number variations (CNVs), and mutational data for patients between TCGA-LUAD and CPTAC-LUAD were obtained from cBioportal (**https://www.cbioportal.org/**).^13^ The cases with the deficiency of tumor staging and overall survival (OS) were excluded. Besides, we deleted three patients in TCGA-LUAD who received neoadjuvant chemotherapy before surgery. Additional statements: To assess the specificity and robustness, GSE37745, GSE30219 and GSE19188 were collected.^10,14^ In addition, we validated transcriptional factors (TFs) activity using GSE148071.^15^ Detailed information about the patients in this study is shown in **Supplementary Table 1**,**2**.

### 2.2 Transcriptional cluster distribution and pathway analysis

For differentiating clusters, a 42-gene classifier was used to identify three transcriptional clusters in all expression datasets through NMF (**Supplementary Table 3**).^16^ Note that hierarchical clustering (HC) was used as orthogonal verification. For HC, median normalized expression and complete distance were adopted. A high confidence queue is considered to have an overall accuracy of over 0.8 and Kappa over 0.6 after cross-validation. Additionally, one thousand highly variable gene expression matrix was used as input for principal component analysis using the R packages “**factoextra**” and “**FactoMineR**” to assess the dissimilarity of the clusters.

Using the **limma** package of R, differential expression gene (DEG) analysis was performed.^17^ DEG was defined as log fold-change > 0.7. To characterize the diversity of clusters, we selected metagenes from previous publications (**Supplementary Table 3**).^3,6,18^ Above scores were calculated on each sample using the R package **GSVA**.^19^

### 2.3 Immune cell evaluation

For previously complete immune-related gene sets, gene expression matrices of Pearson’s correlation (R_min_ > 0.5) were used to select the streamlined version in treatment-naive and ICIs-treated NSCLC cohorts.^18,10,20,21,22^ The higher correlation to elucidate reproducible patterns of gene expression, the more relevant to the clinic. Finally, effector cells (effector memory CD8^+^ T and T helper 1 cells), and immunosuppressive cells (Tregs and MDSCs) were retained from the 28 gene sets, which are highly tumor-specific (**Supplementary Table 3**).^18^

For immune cell infiltration, the CIBERSORT website (**http://cibersort.stanford.edu**) was used to repeat 1000 times to assessment for the relative infiltration proportion of 22 types of immune cells.^23^ Patients with a p-value less than 0.05 were retained. Also, tumor purity was evaluated by the ESTIMATE method.^24^

### 2.4 Genetic perturbation

For methylation analysis, we downloaded methylation data in the Xena website.^12^ We evaluated the signal value of MHC-II enhancers. MHC-II enhancers revealed to be overlapped with lung-specific enhancers from Mullen et al.^25^ The intensity of methylation was quantified by the β value (0-1) of each CpG site. Moreover, we collected global methylation information for patients from Jung et al.^22^

To evaluate super-enhancer activities, we obtained the expression matrix of enhancer RNA (eRNA) from **https://bioinformatics.mdanderson.org/Supplements/Super_Enhancer/5_Super_enhancer_annotation/TCGA_RPKM_eRNA_300k_peaks_in_Super_enhancer_LUAD.txt.gz**.^26^ For assessment of the global activation of super-enhancers, global eRNA level was estimated through two thousand highly variable eRNAs. Furthermore, we performed the R package **ImmuLncRNA** to determine immune-specific eRNA.^27^

### 2.5 Regulation network

For single-cell RNA (scRNA) sequencing, we downloaded the GSE148071 raw expression matrix and conducted downstream analysis by **Seurat** R package.^15,28^ To infer the activity of TFs, we randomly sampled 1000 cells from GSE148071 and stratified them by the median value of the immune signature.^15,18^ Notably, cisTarget databases storing information about TF were downloaded according to the default settings.

Then, the flow-sorted epithelial cell profile (EPCAM^+^ CD45^-^ CD31^-^) was downloaded and converted into a regulator matrix using the R package **dorothea**.^29,30^ The R package **dorothea** relies on the VIPER algorithm and incorporates experimental evidence intending to predict TF’s activity by RNA datasets.

### 2.6 Preclinical models utilization

CMAP database (**https://clue.io**) which stored predicted compounds perturbation was used.^31^ Optional query functions for the CMAP database include gene expression, cell viability, and proteomics. Gene expression and cell viability functions were used in this study, requiring the input of gene signatures and cell line names. The positive value of compounds reflects a consistent trend with the phenotype, but not negative values. To obtain high-quality compounds, we only considered predictions with scores greater than absolute 1.5.

The cell lines RNA sequencing matrix was from Broad Institute (**https://depmap.org/portal/**).32 Only 127 NSCLC CCLE cell lines labeled as “type-refined==NSCLC” were selected (HCC1588 was excluded because it was associated with COAD). NSCLC UTSW cell lines for validation from McMillan et al.33 Beyond, PDX RNA sequencing data of NSCLC were from NCI-MATCH trial and Asian cohort GSE78806.^34,35^ All NMF-identified preclinical models including cell lines and PDXs were described in **Supplementary Table 4**.

For drug repositioning, CTRP AUC data were from **https://ocg.cancer.gov/programs/ctd2/data-portal**.^32^ Meanwhile, cell lines expression and IC50 matrix from GDSC were downloaded in **https://www.cancerrxgene.org/**.^36^ AUC and IC50 values were defined as a measure of drug sensitivity. We generated per-drug sensitivity scores for each sample via the R package **oncoPredict**.^37^ The **oncoPredict** R package was used to predict the relative sensitivity of monotherapy based on a batch-corrected expression profile. In addition, we use the manuscript’s orthogonal discovery method, called a random forest, which can effectively capture potential gene profiles regarding drug sensitivity.^38^

### 2.7 Cell culture and cell viability

The lung adenocarcinoma H1944 cell line was purchased from Pricella, while the BEAS-2B cell line was obtained from the Xinqiao Hospital Cancer Institute. H1944 and BEAS-2B were maintained in RPMI 1640 (SH30809.01 cytiva) supplemented with 10% FBS and 1% streptomycin in 5% CO2 at 37 °C. Cell viability was estimated by CCK-8 assay. BEAS-2B and H1944 cell lines were co-cultured with Dinaciclib (HY-10492, MedChemExpress) for 48 and 72 hours, respectively. Meanwhile, treatment of the H1944 cell line were with Alvocidib (HY-10005, MedChemExpress) for 72 hours.

### 2.8 Western blot analysis

In brief, the H1944 cell line was treated with Verteporfin (HY-B0146, MedChemExpress) for 96 hours. Then, RIPA buffer (P0013B, Beyotime Biotechnology) containing 1% PMSF (ST2573, Beyotime Biotechnology) was added to the cell culture for lysis. Cell lysates were collected as supernatants after centrifugation and the total protein content was determined by the BCA method. Equal proteins (20 μg/lane) were loaded on SDS-PAGE gels and then transferred to pvdf membranes. After 1 hour of closure with Western closure solution (BL535A, Biosharp), pvdf membranes incubate with the primary antibody overnight at 4 °C. Next, the membrane was incubated with the secondary antibody for 1 hour at room temperature after washing with TBST. Finally, the target bands were visualized using a chemiluminescent imaging system (FluorQuant AC600, AcuronBio). The antibodies used include rabbit anti-KLF5 polyclonal antibody (Cell Signaling Technology), and rabbit anti-β-actin polyclonal antibody (BL005B, Biosharp). All bands were normalized to β-actin and the bands were analyzed by Image J.

### 2.9 Statistical Analysis

The R package **survminer** was used to plot the OS by the Kaplan-Meier analysis. Heatmaps were based on Z-value normalized gene expression, as previously described. Pearson’s chi-squared test or Fisher’s exact test was applied to compare all proportions. The non-parametric test (Wilcoxon-test or Kruskal-Wallis test) was performed to compare all variables between groups. All analytical tests were two-sided. A value of p < 0.05 was considered to be statistically significant. All codes used for analyses were written in R software.

## 3. Results

### 3.1 Transcriptional expression profile classifies three prognostic clusters

We assume that the smallest subset will benefit from targeted therapies. According to our observations, the most pronounced neuroendocrine profile was in the PP-3 subtype, which may reflect the original PP subtype (**Supplementary Table 5**).^39^ Indeed, the NMF algorithm can decompose the expression matrix into small metagenes.^16^ Using the NMF method, a cohort of three hundred and twenty surgically resected stage IB–IIIA tumors from TCGA-LUAD was designed with three clusters after overall consideration (**Fig. S1A-B**). The 42-gene classified patients as bronchioid, neuroendocrine and squamoid clusters (**Fig. S1C**). As expected, the PP-3 subtype is a subset of our neuroendocrine cluster (**Fig. S2A**).^39^ It is interesting to note that the mesenchymal subtype is also a subset of the bronchioid cluster (**Fig. S2B**).^40^ Although moderately similar to our cluster in TCGA-LUAD, not related in Soltis et al (**Fig. S2C, Supplementary Table 2**).^5,41^ Despite the concept of three lung lineage being widely accepted, we believe it is necessary to consider the cohort’s inclusion criteria. Thus, an orthogonal approach called HC is used to obtain high-confidence queues including CPTAC-LUAD, GSE72094, and TCGA-LUAD (**see “methods”**) (**Supplementary Table 6**).

Furthermore, we first explore prognostic value among three clusters, the shortest OS was in the squamous cluster (**Fig. 1A-B**). Then univariate analyses show that age, tumor staging, and squamous cluster were significantly prognostic (**Supplementary Table 7A**). After adding age, tumor staging, and gender, squamous is still an independent prognostic factor using multivariate Cox analyses. Meanwhile, we found a moderate association of clusters with clinical characteristics (**Supplementary Table 7B**). After we demonstrated the prognostic significance of the cluster, we asked whether there were significant differences between clusters. Principal component analysis analysis showed that three acquired clusters could be divided with highly variable gene expression data (**see “methods”**) (**Fig. 1A-B**). A previous study revealed that PP and PI subtypes had similar expression distribution of NKX2-1.^42^ Among three clusters, we noticed the squamoid cluster was highly overlapping with patients with lower quarter expression of NKX2-1 (over 90%, **Fig. S1D-E**). Moreover, clusters could distinguish the different fates among three cohorts (**Supplementary Table 7C**), supported for previous descriptions.^7^ In addition, proposed clusters overcome the challenges of RNA-Protein concordance and tumor heterogeneity, and are confirmed in samples and PDXs (**Supplementary Table 2**,**4**). To validate the clinical value of clusters, we examine the prognostic significance in patients with adjuvant chemotherapy. Our results showed that the squamoid cluster had an improved prognosis after receiving cisplatin-based therapies, and the bronchioid cluster possessed potential efficiency with pemetrexed monotherapy (**Fig. 1C-D**).

**Figure 1.**
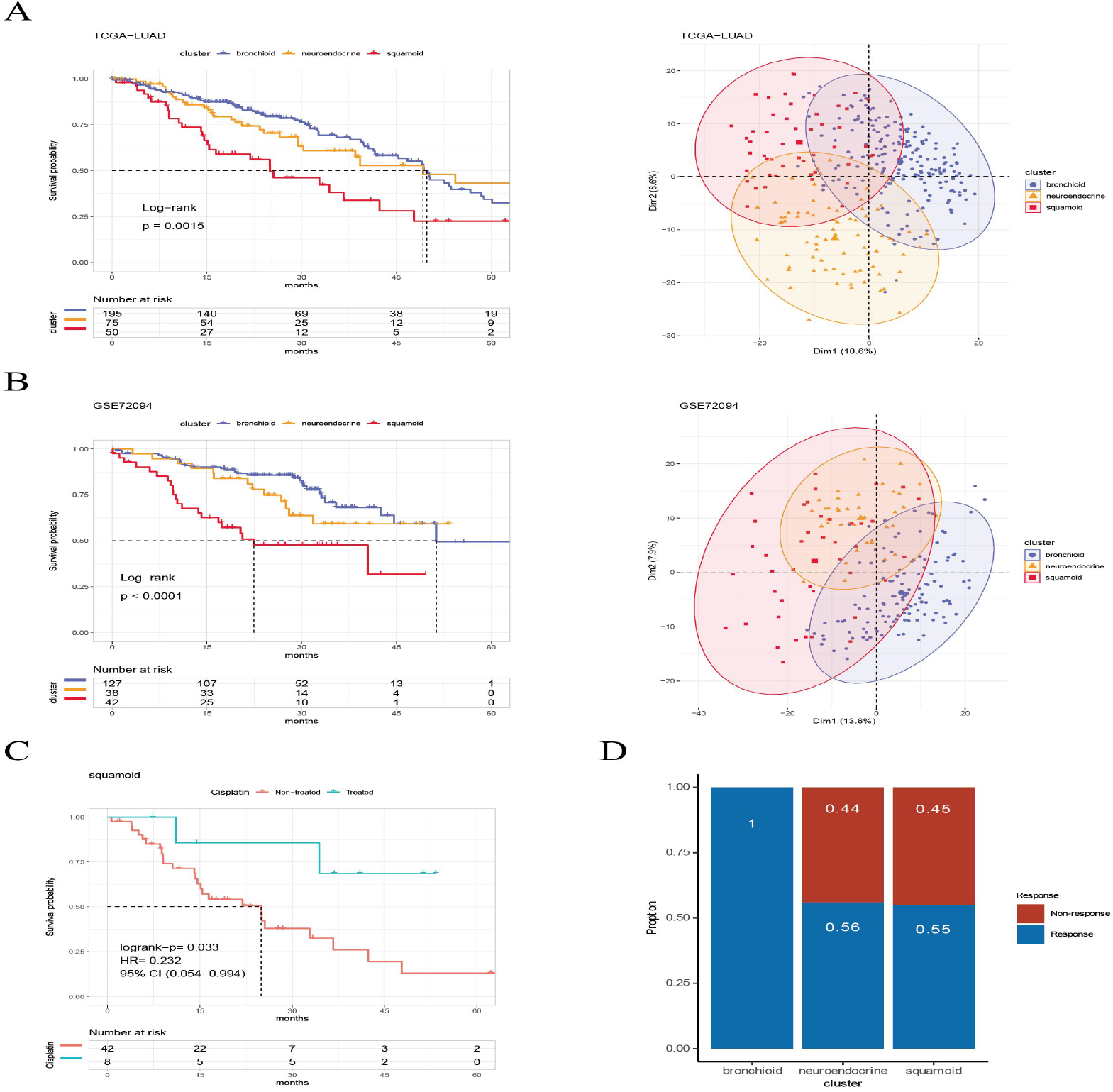
Prognosis and predictive chemotherapy information of transcriptional clusters. Kaplan-Meier Survival is showing prognostic significance by Log-rank test and PCA analysis plotting patients were clustered into three distinct clusters for (A) TCGA-LUAD cohort (n = 320), and (B) GSE72094 cohort (n = 207) (left: survival analysis; right: PCA plotting, the X and Y axes represent the variability in the differentiated clusters of Dim1 and Dim2, respectively). (C) Squamoid clusters received cisplatin alone or combined therapy from TCGA-LUAD (n = 50) and (D) GSE19188 (n = 70) predicted for pemetrexed therapy. Legend was labeled in blue (bronchioid), yellow (neuroendocrine) and red (squamoid).

### 3.2 Comprehensive characterization among three clusters

We selected four lung-specific metagenes to represent clusters described in **Supplementary Table 3**.^3,6,18^ Using the R package **GSVA**, we found that neuroendocrine and squamous shared proliferation among four cohorts (**Fig. 2A**).^19^ Although metagene patterns were shared in different clusters, the metagenes of surfactant, neurodevelopment, and basal distinctly corresponded to bronchioid, neuroendocrine, and squamoid, respectively, suggesting that metagenes can reflect approximate transdifferentiation directions among clusters. We hoped to understand the immune profiles in transdifferentiation-related clusters. Among three cohorts, the least immune infiltrate was in the neuroendocrine cluster (**Fig. 2A**). The bronchioid cluster had the highest proportion of resting mast cells, but the lowest proportion of activated memory CD4^+^ T cells using CIBERSORT algorithm (**Fig. 2B**).^23^ After considering high-resolution dataset, we found squamous cluster was most related to T-cell status but had the least CellPhoneDB inferred cellular communication using Scissor method (**Fig. S3A-C, Supplementary Table 8**).^43,44^ This may be related to the solid pattern driven by negative NKX2-1.^45^

**Figure 2.**
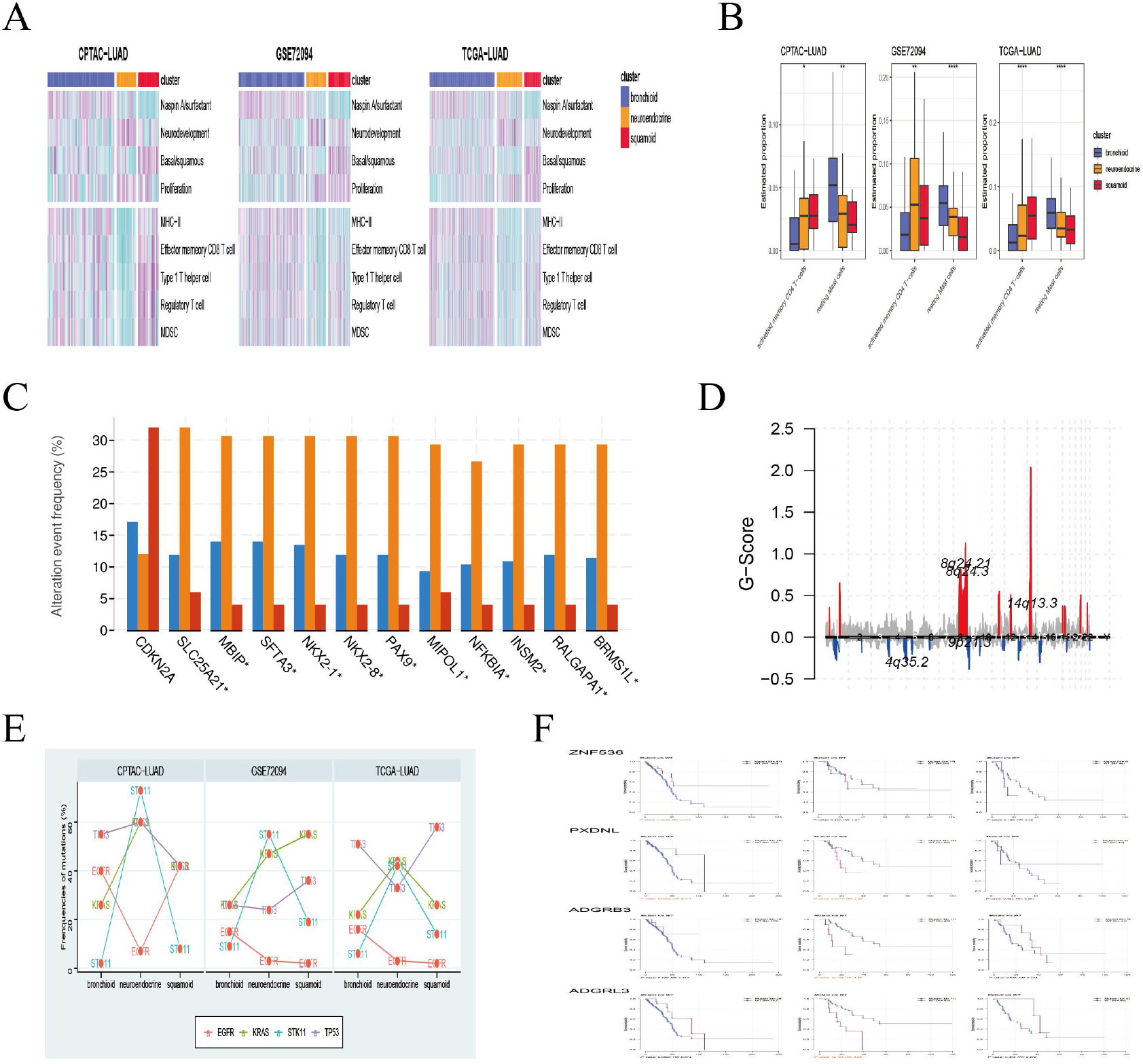
Pathways, immune cells, copy number aberrations and mutations among three clusters. (A) Heatmap drawing Gene Set Variance Analysis (GSVA) scores for each patient in the CPTAC-LUAD cohort (n = 77), GSE72094 cohort (n = 207) and TCGA-LUAD cohort (n = 320). Z-value GSVA score projected into (−2;2). (B) Box plots exhibiting comparison of immune cell evaluated proportion using Kruskal-Wallis test in the CPTAC-LUAD cohort (n = 77), GSE72094 cohort (n = 207) and TCGA-LUAD cohort (n = 320). *, **, *** and **** representing p<0.05, p<0.01, p<0.001, respectively. (C) CDKN2A deletion and 14q13.3 amplifications among three clusters in the TCGA-LUAD cohort. (D) Genome plot showing focal chromosomal alterations of neuroendocrine in the TCGA-LUAD cohort. Given the G-score (x axis) for each focus events (y axis). Note that high G-score means high probability of occurring event. (E) Line graph showing the percentage distribution of the four major mutations (EGFR, KRAS, STK11 and TP53) between clusters in the CPTAC-LUAD, GSE72094 and TCGA-LUAD cohorts. (F) Kaplan-Meier plot showing survival time of ZNF536, PXDNL, ADGRB3 and ADGRL3 mutation status among three clusters in the TCGA-LUAD cohort (left: bronchioid; middle: neuroendocrine; right: squamoid).

Next, we examined the CNVs among three clusters, many 14q13.3 features (e.g., NKX2-1 and MBIP) were enriched in the neuroendocrine cluster.^46^ While the squamous subtype had the highest CDKN2A variation but the lowest 14q13.3 (**Fig. 2C**). Further examination revealed that the neuroendocrine cluster had a high frequency of chromosome 8q variation, especially 14q13.3 using the R package **maftools** (**Fig. 2D**).^47^ Furthermore, we focused on four major mutations (EGFR, KRAS, STK11, and TP53) in CPTAC-LUAD, GSE72094 and TCGA-LUAD cohorts (**Fig. 2E**). Importantly, STK11 mutations were mainly distributed in the neuroendocrine cluster. The distribution of mutations among clusters is very different across three cohorts (e.g., KRAS and EGFR), which may be due to ethnicity and smoking history. In conclusion, the neuroendocrine cluster is characterized as an immune cold partially because of STK11 mutations, which show resistance to ICI therapies (**Fig. 2A, 2E**).^1,2^ Then, mutations with prognostic significance in clusters were paid attention to ZNF536, PXDNL, ADGRB3, and ADGRL3 mutations were almost greater than 10% in frequency by in the cBioportal website (**Fig. S4A**). In the bronchioid cluster, the mutant type of ZNF536 and PXDNL showed longer OS than the wild type, the other clusters were the opposite (**Fig. 2F**). Similarly, ADGRB3 and ADGRL3 are likely oncogenic drivers in the neuroendocrine cluster, while they exert tumor suppressors’ role in other clusters. We further explored whether the above mutations were relevant to ICI therapies. Among these four mutations, only ZNF536 mutations showed a better prognostic trend in both cohorts (**Fig. S4B**). Meanwhile, ADGRB3 mutations are related to a favorable prognosis in Hellmann’s nonsquamous NSCLC cohort receiving PD-1 plus CTLA-4 blockade (**Fig. S4C**). Overall, genomic results are presented among clusters, and the neuroendocrine cluster has multiple genetically altered vulnerabilities, such as STK11 mutations and NKX2-1 amplification.

### 3.3 Gene classifier dissects genetic perturbations involved in immunity

Considering that low expression of major histocompatibility complex II (MHC-II) is a distinctive feature of the neuroendocrine cluster, we gain further insight into the underlying mechanisms (**Fig. 2A**). A previous study suggested the expression of MHC-II was mainly regulated by enhancers, the bronchial, squamous and neuroendocrine clusters from highest to lowest according to the level of MHC-II enhancers (**Fig. 3A**).^21^ We also found a similar trend in global-methylation level (**Fig. 3B**). Importantly, there was a high correlation coefficient (R = 0.47, **Fig. 3C**) between MHC-II and global-methylation level, supported by the previous study.^22^ These results imply that perturbations in genetics can affect TME among three clusters.

**Figure 3.**
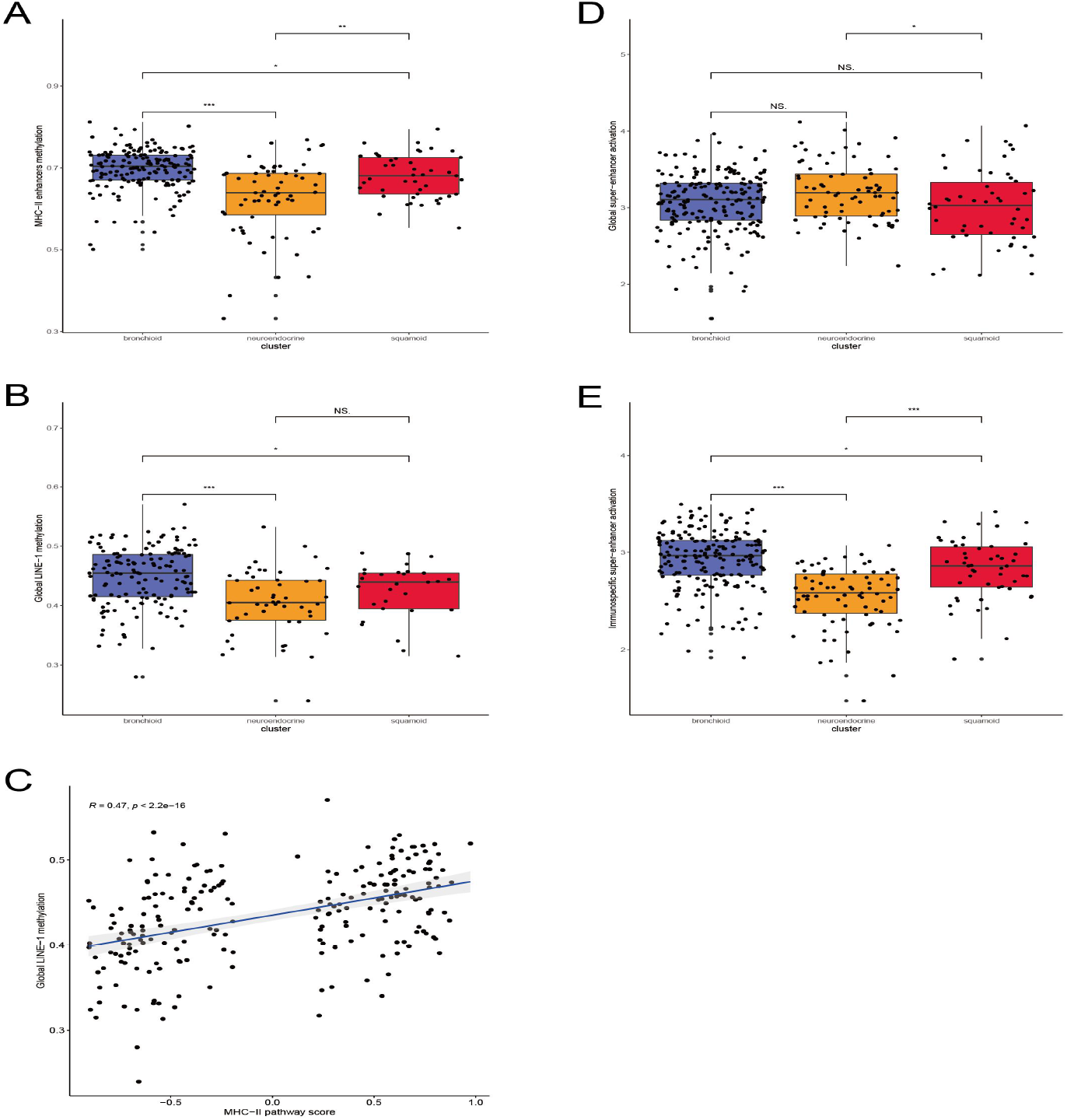
Methylated and super-enhancers perturbations between clusters. Box plots showing the levels of methylation from (A) MHC-II enhancers and (B) Global LINE-1. (C) Spearman’s correlation coefficient analysis between the level of MHC-II and Global LINE-1. Box plots exhibiting the expression from (D) Global super-enhancer activation and (E) Immunospecific super-enhancer activation in the TCGA-LUAD cohort. Comparison between groups using Wilcoxon-test. *, **, *** and **** representing p<0.05, p<0.01, p<0.001, respectively.

Next, we want to explore whether genetic perturbations are specific to the immune activity. The eRNA can predict enhancer activities and reflect uncontaminated signals compared with RNA.^26^ In this study, we focused on super-enhancers, which were described as large regions with enhancer activity. By quantifying the two thousand highest variable eRNAs, the highest expression level was found in the neuroendocrine cluster (**Fig. 3D**). Interestingly, the opposite result occurred between the clusters when using immuno-specific eRNAs (**Fig. 3E**). In conclusion, our results suggest that the bronchioid cluster shows the highest level of immune activity in three aspects: immuno-specific super-enhancer, MHC-II enhancers, and global LINE-1 methylation.

### 3.4 Transcriptional clusters design personalized targeted therapy

To develop combined targeted chemotherapy or targeted immunotherapy, we first used DEGs among the three clusters and back-validated with NMF-annotated cell lines in the CMAP database (**Supplementary Table 9**).^31^ Despite no common drugs being identified, multiple MEK inhibitors may be resistant in the bronchioid cluster, which may be contrary to what was reported by Daemen et al (**Fig. S5A**).^40^ Interestingly, DEGs of subtypes may be druggable targets, such as SLC34A2 (**Supplementary Table 9**).^48^ We further analyzed the CCLE and GDSC databases, and for GDSC only the top 100 compounds with high lethality were included (**Supplementary Table 9**). Through larger-scale drug-sensitive datasets predicted by R package **oncoPredict**, our results show that MEK inhibitors may favor squamoid cluster based on CCLE and GDSC datasets (**Fig. S5B-C**).^37,32,36^ Specifically, the integration of patient information with cell lines, which is not available in the CMAP database.

Available evidence suggests neuroendocrine clusters are consensus, we found that CDK inhibitors including dinaciclib and alvocidib have preferred activity in the neuroendocrine cluster (**Fig. 4A**).^5,40,49^ As expected, alvocidib could inhibit the proliferation of NCI-H1944, which was identified as a neuroendocrine cluster (**Fig. 4B**). Given that the second-generation pan-CDK inhibitor dinaciclib may be superior to than the first-generation pan-CDK inhibitor alvocidib, we further validated the proliferative effect of dinaciclib in tumor and normal epithelial cell lines (**Fig. 4C**). Our results suggest that dinaciclib is promising in the neuroendocrine cluster and that we should be aware of concentration.

**Figure 4.**
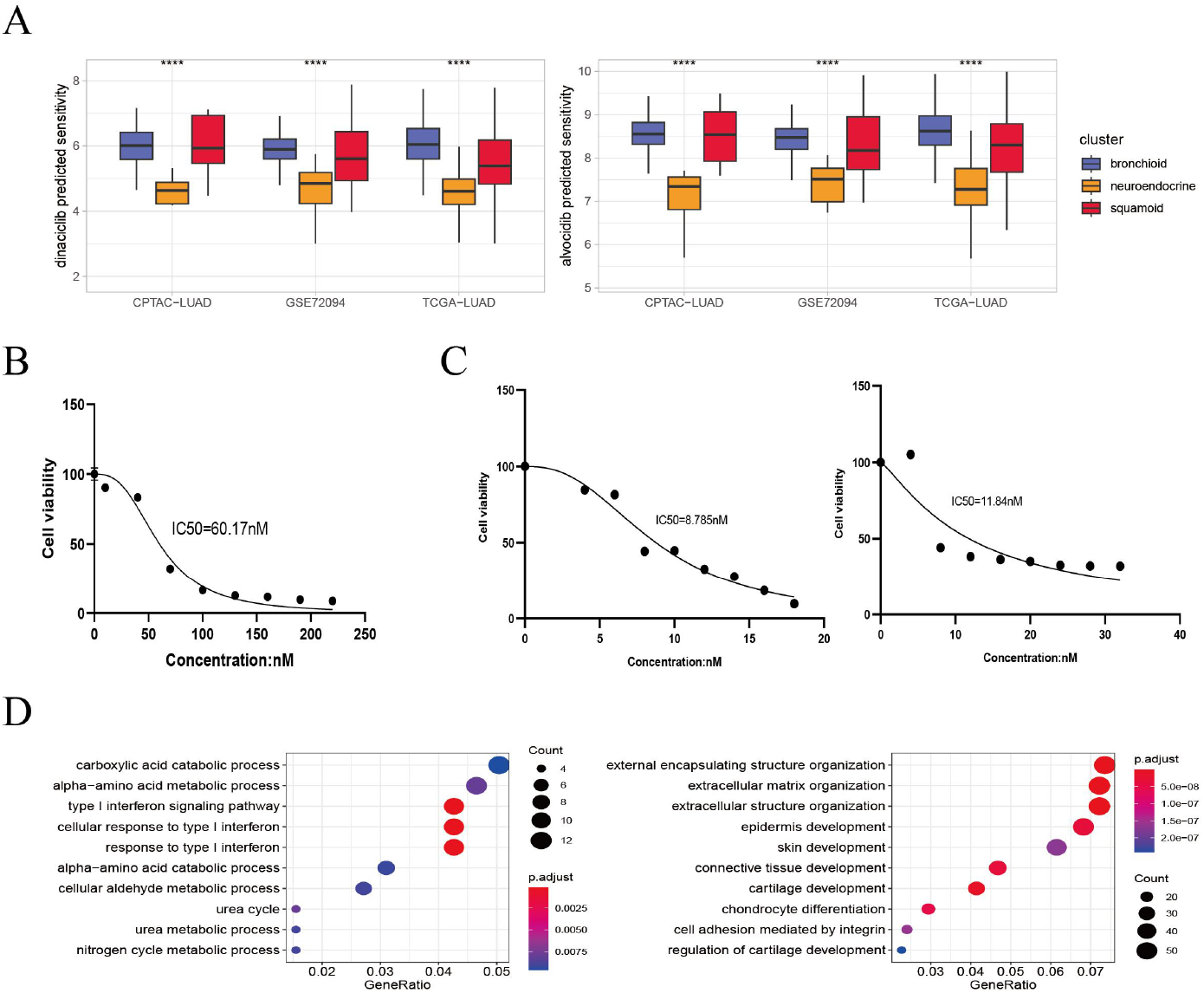
Dinaciclib and alvocidib predicted from bioinformatics and validated in vitro. (A) CCLE predicted sensitivity values in clusters using Kruskal-Wallis test. *, **, *** and **** representing p<0.05, p<0.01, p<0.001, respectively (left: dinaciclib; right: alvocidib). (B) Cell viability of the NCI-H1944 cell line treated with alvocidib. (C) Cell viability of the NCI-H1944 and BEAS-2B cell lines treated with dinaciclib (left: NCI-H1944; right: BEAS-2B. The horizontal and vertical axes are the concentration and cell survival ratio, respectively, while the IC50 values have been labeled). (D) GO enrichment analysis of NCI-H1944 after treatment with dinaciclib and alvocidib, respectively (left: down-regulated shared pathways; right: up-regulated shared pathways).

A recent study proved drug sensitivity could be inferred through expression profiles.^38^ We asked whether one nonlinear regression model named random forest could discover drugs, with similar functional mechanisms. Interestingly, PLK1 inhibitors and EGFR inhibitors show a high degree of similarity (about 20%-50%). In addition, both dinaciclib and alvocidib reduced interferon and metabolic processes, and increase extracellular matrix and squamous remodeling (**Fig. 4D**). Taken together, our results validate for the first time that dinaciclib and alvocidib are highly similar (model prediction: about 50%; transcriptome results: alvocidib: 79%, dinaciclib: 43%, **Supplementary Table 10**,**11**), and imply this algorithm has the potential to identify more similar compounds.

### 3.5 Networks identify KLF5 as a driver of lineage development and immune invasion

Using a large NSCLC-cohort (n = 5589, **Supplementary Table 1**), our previous view was refined, and further, dissect the pre-existing TME specifically into effector and suppressor cells.^18^ We obtained robust immune gene sets and proved that immune cold means complete resistance to PD-L1 blockade, although immune hot does not guarantee benefits. Together with our results, the abundance of Treg cells is also a potential indicator of anti-PD-1/PD-L1 therapies (**Fig. 5A**).^4^

**Figure 5.**
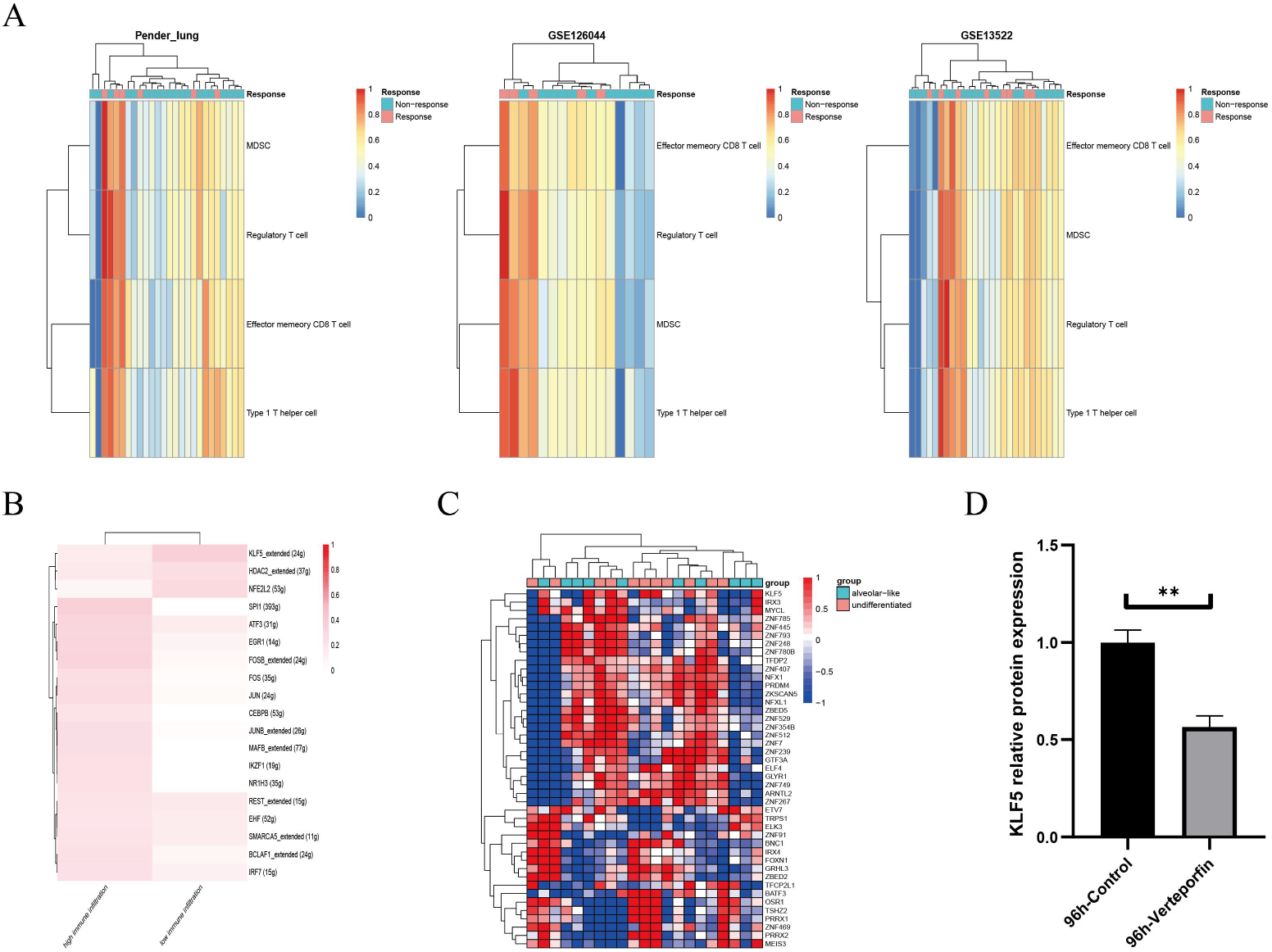
KLF5 as a targetable target inferred from single cells and bulk RNA sequencing. (A) Heatmap plotting the relative immune infiltration based on Gene Set Variance Analysis (GSVA) scores in Pender’s lung (n = 25), GSE126044 (n = 16) and GSE135222 (n = 27) cohorts (left: Pender’s lung; middle: GSE126044; right: GSE135222, Z-value score projected into (0;1)). (B-C) Transcriptional factors regulation grouped by median signature score in GSE148071 and GSE111907, respectively (GSE148071: 0 means no activity, 1 the opposite; GSE111907: Z-value score projected into (−1;1)). (D) Verteporfin-induce alterations in the protein level of KLF5 in the NCI-H1944 cell line. The western blot analysis-derived bands were normalized to β-actin.

We assume that lung lineage development maintains immune balance. Two networks were constructed based on scRNA and epithelial bulk sequencing, using the immune and epithelial gene sets, respectively (**Fig. 5B-C**).^15,29,18,50^ Lower immune infiltration may be associated with aggressive tumor progression. Our results suggest that KLF5 inhibited immune activation and bronchial differentiation. Given KLF5 may be recruited when YAP1 phosphorylation, we examined whether verteporfin could inhibit KLF5.^51^ As expected, verteporfin inhibited protein levels of KLF5 (**Fig. 5D**). We speculate that verteporfin regulates the TME partly through the lineage factor KLF5. Future studies to determine whether verteporfin resolves KLF5-dependence would be interesting.

### 3.6 Clinical translation of transcriptional clusters

We first assessed common clinical indexes related to immunotherapy including TMB, CNVs, and tumor purity.^1,2^ Among the three clusters, the neuroendocrine cluster had highest TMB, CNVs and tumor purity (**Fig. S6A-B**). Both tumor purity and cross-platform influenced the robustness of clusters. We believe that the single sample predictor algorithm generated by our clusters is promising for testing personalized therapies through NanoString nCounter technology.^52^

Our cluster not only reflected local perturbation-related immunity but also complemented the existing immunophenotypes.^53^ We observed the proportion of immunophenotypes finding large inconsistencies in different datasets (e.g. TGF beta dominant). Nonetheless, the relative distribution between clusters and immunophenotypes was evident, as reflected in the bronchioid cluster, mainly in lymphocyte depleted (**Fig. S7**). Overall, transcriptional clusters could provide additional information for immunotherapy. A study showed an approximate 63% overlap between NKX2-1 and MHC-II in positive immunohistochemistry results.^54^ Thus it is reasonable to assume that NKX2-1 promotes anti-tumor immunity and sensitivity to immunotherapy by stabilizing MHC-II.^54,55^ Recently ORIENT-11 trial showed that high expression of MHC-II is associated with the TME that responds to combination therapies.^3^ Combined with our results, subtypes may reveal therapeutic vulnerabilities. Hence, we propose a molecular hypothesis to facilitate clinical practice (**Fig. 6**).

**Figure 6.**
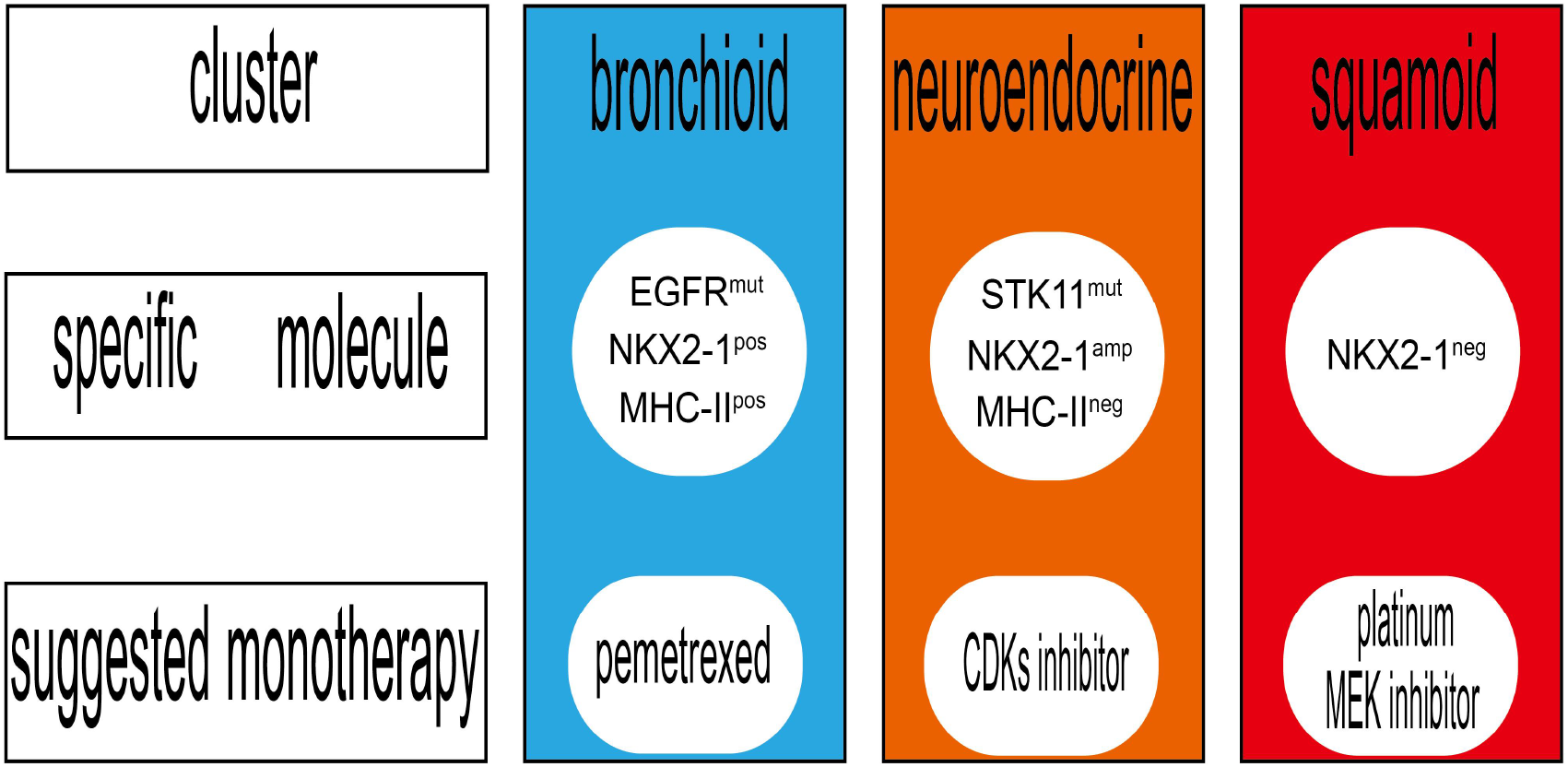
Graphical abstract. The viable therapeutic strategies among bronchioid, neuroendocrine and squamoid.

## 4. Discussion

In this study, we established three clusters across samples, PDXs, and cell lines. Prognosis, genomics, genetics, and TME were depicted among bronchial, neuroendocrine and squamoid clusters. We proved that immune pathways did recognize lineage factors, and focused on the analysis of drug sensitivity differences in three lung cancer lineages.

Our results confirmed that classical pathways can profile molecular characteristics of clusters. The bronchioid cluster had high surfactant levels (e.g., NAPSA and SFTPC, etc.), and the neuroendocrine cluster showed the activated-neurodevelopment (e.g., ASCL1 and INSM1, etc.), and high expression of keratin genes (e.g., KRT6A and KRT16, etc.) was shown in the squamous cluster. Among the three clusters, the bronchioid cluster is associated with non-smokers and females, while neuroendocrine and squamous clusters in contrary. Recently, Roh et al. depicted vulnerabilities in PI subtype, whereas we tended to refine precise PP subtype.^49,39^ The neuroendocrine cluster was defined as having the highest frequency of STK11 mutations and NKX2-1 amplifications, but the lowest immune infiltration. It is confirmed that bronchioid is immune activated, which may be due to EGFR mutations neglected in the past.^1,2^ Indeed, NKX2-1 may be contradictory in immunotherapy, with NKX2-1 positive individuals having higher MHC-II, while immunosuppressive TME in negative NKX2-1 patients. We speculate that the squamoid cluster may have the worst prognosis due to the hybrid state.^56^

From the therapeutic aspect, the squamoid cluster may benefit from the MEK inhibitors after large-scale drug sensitivity analysis. Recently described dysregulation of CDKs in the neuroendocrine cluster, we further show that patients with high proliferation but low squamous and extracellular matrix remodeling may benefit from pan-CDK inhibitors.^41,49^ Combined with our results, dinaciclib is the potential to be repositioned, considering precise subset and reduced toxicity.^57^ Importantly, our clusters may reflect NKX2-1 biology.^7,42,45,46,54,55^ The CMAP database provided not only compounds but also knockout or overexpression gene information, which may help phenotype-based drug repositioning.^31^ Another concept is network-based drug repositioning, we constructed an immunotherapy-related scRNA regulatory network and recognized potential regulators by epithelial bulk sequencing. Interestingly, a lineage factor named KLF5 stood out from the above two networks, and verteporfin did inhibit the level of KLF5. In fact, verteporfin could bind to KLF5 protein via AutoDock software prediction (data not shown).

Emerging evidence suggests that scRNA analysis was good at resolving heterogeneity, but had difficulty in simultaneously identifying cancerous and non-cancerous components.^56,58^ In this study, we implemented two alternative strategies, one focusing on scRNA integration bulk and the other on flow-sorted epithelial profile. Li et al. showed co-expression of lineage, however, bronchioid became overwhelmingly dominant and did not determine the lineage classification of patients.^56^ In contrast, the transcriptional clusters allow for personalized therapies in identifiable lung cancer lineage.^7,8,9^ Additionally, we compare correlation analysis of global, lineage-specific, our 42-classifier and immune pathways genes, separately (**Fig. S8A-B**). Lower correlations from cell lines were observed considering lineage and our 42-classifier. Moreover, about 86% of lineage-specific genes were a low expression in McMillan et al.^33^ Our results show that partly lineage features of the cell lines were lost, thus the validity of preclinical models needs to be considered.^7^

The direction of future research is the combination of genetic and non-genetic, and arguably epistatic oncogenic transcriptomic landscape. Our study focuses on the design of non-genetic aspects, the important questions are whether there is a need to intervene in transdifferentiation and how to rely on the lineage to target therapeutic vulnerabilities. Subtypes may be confounded by clinical factors, such as smoking. We observed all neuroendocrine belongs be smokers in the CHOICE cohort, thus the absence in the OncoSG cohort is explainable (**Supplementary Table 2**).^59,60^ To our knowledge, the TRU and non-TRU binary classification may apply to cohorts with a high proportion of non-smokers.^52^ Despite these shortcomings, we propose RNA-Protein consistent clusters and focus on transcriptome-based drug repositioning, and in the future, we will also integrate chromatin accessibility and radiomics. Also, implement of metagenes via the NanoString platform is a value.

## Ethics approval and consent to participate

Not applicable

## Consent for publication

Not applicable

## Methods and availability of supporting data

All data generated and methods described were in accordance with the relevant guidelines and are permitted by non-commercial organization and did not need access approval. The corresponding author can be contacted for reasonable data.

## Competing interests

Not applicable

## Funding

This study was supported by the National Natural Science Foundation of China (81773245, 81972858, 82202951 and 82172670), the Technology Innovation and Application Development Project of Chongqing (cstccxljrc201910) and the Cultivation Program for Clinical Research Talents of Army Medical University (2018XLC1010 and 2019XQN10).

## Contributions

J. S. designed the study. L. Zeng and L. Li collected data. L. Zeng and L. Zhang performed analyses. X. Liao, C. Yin, X. Chen and L. Zhang wrote the text. All authors reviewed the manuscript.

## Acknowledgements

We thank BGI Genomics Co., Ltd (Shenzhen, China) for transcriptional sequencing, and appreciate the data from the TCGA, CPTAC and GEO datasets.

## Notes

### Competing Interest Statement

The authors have declared no competing interest.

