## Supplementary figure for "RNA sequencing identifies lung cancer lineage and facilitates drug repositioning"

**SUPPLEMENTARY INFORMATION**

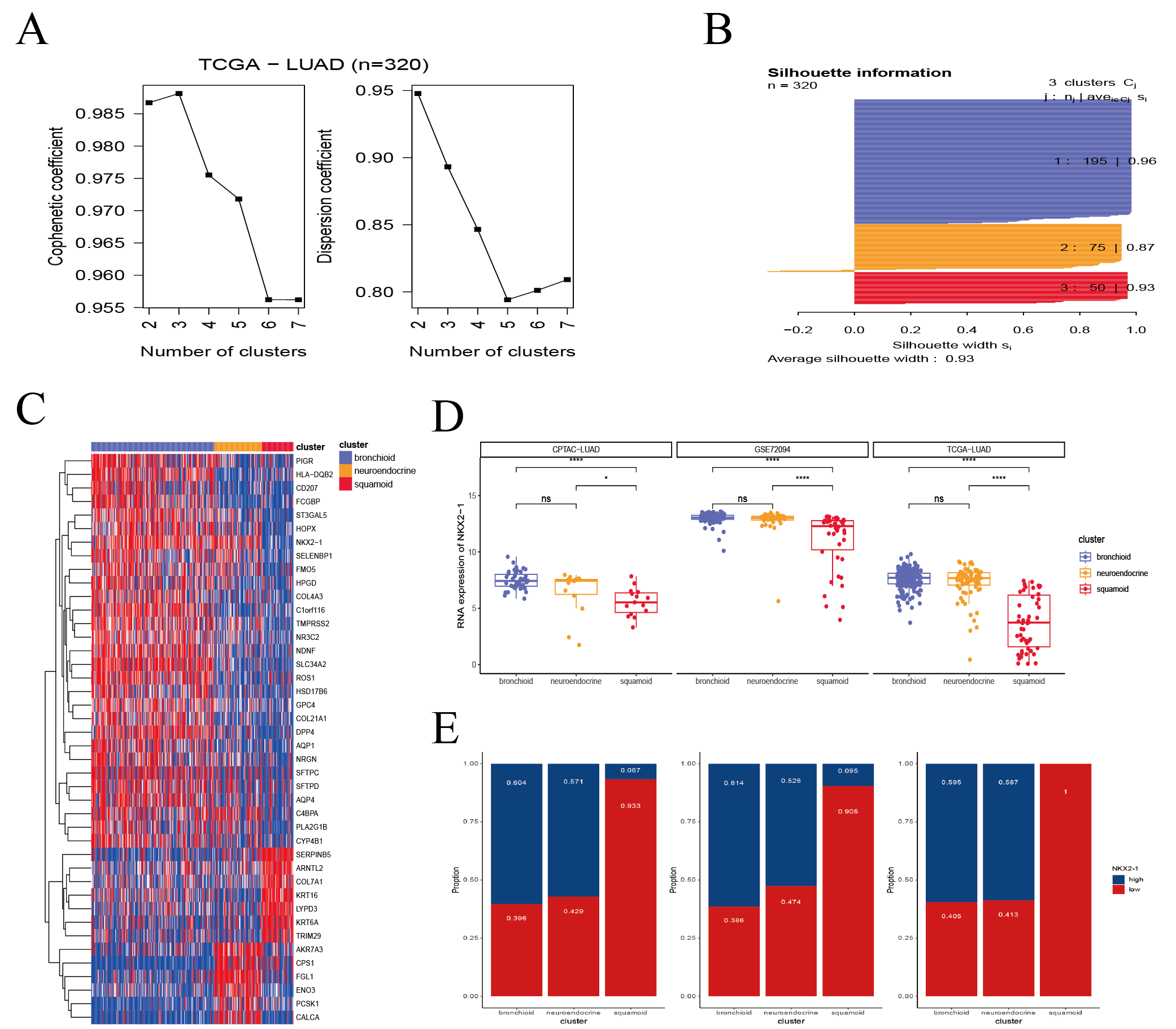

**Figure S1. Clustering analysis classified three prognostic clusters**

(A) Non-negative matrix factorization was performed in the stage IB–IIIA TCGA-LUAD cohort. Cophenetic coefficient and dispersion coefficient were shown for k = 2 to 7. (B) Silhouette information of stage IB–IIIA TCGA-LUAD for k = 3 was exhibited. (C) Heatmap showing two clusters using 42 candidate genes. The expression data was based on Z-value normalization. Z-value score projected into (-1;1). (D) Box plots showing the RNA expression of NKX2-1 among bronchioid, neuroendocrine and squamoid clusters. Comparison between groups using Wilcoxon-test. *, **, *** and **** representing p<0.05, p<0.01, p<0.001, respectively. (E) Bar plots represent the relationship between median defined NKX2-1 group and clusters.

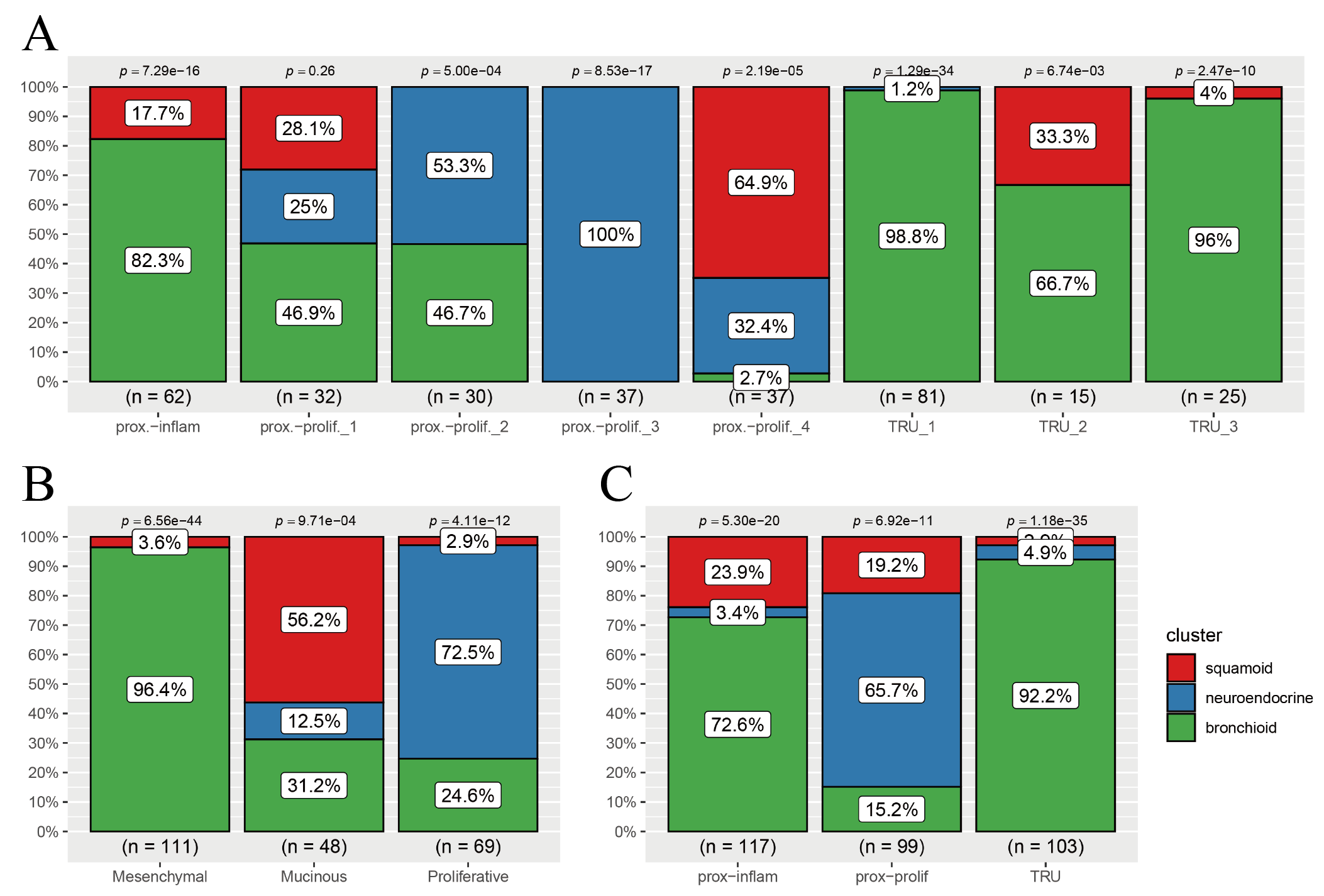

**Figure S2. Relationship between previous subtypes and three clusters**

Bar plots showing overlapping ratio between our clusters and (A) Wang et al, (B) Daemen et al, and (C) TCGA group subtypes in the TCGA-LUAD cohort.

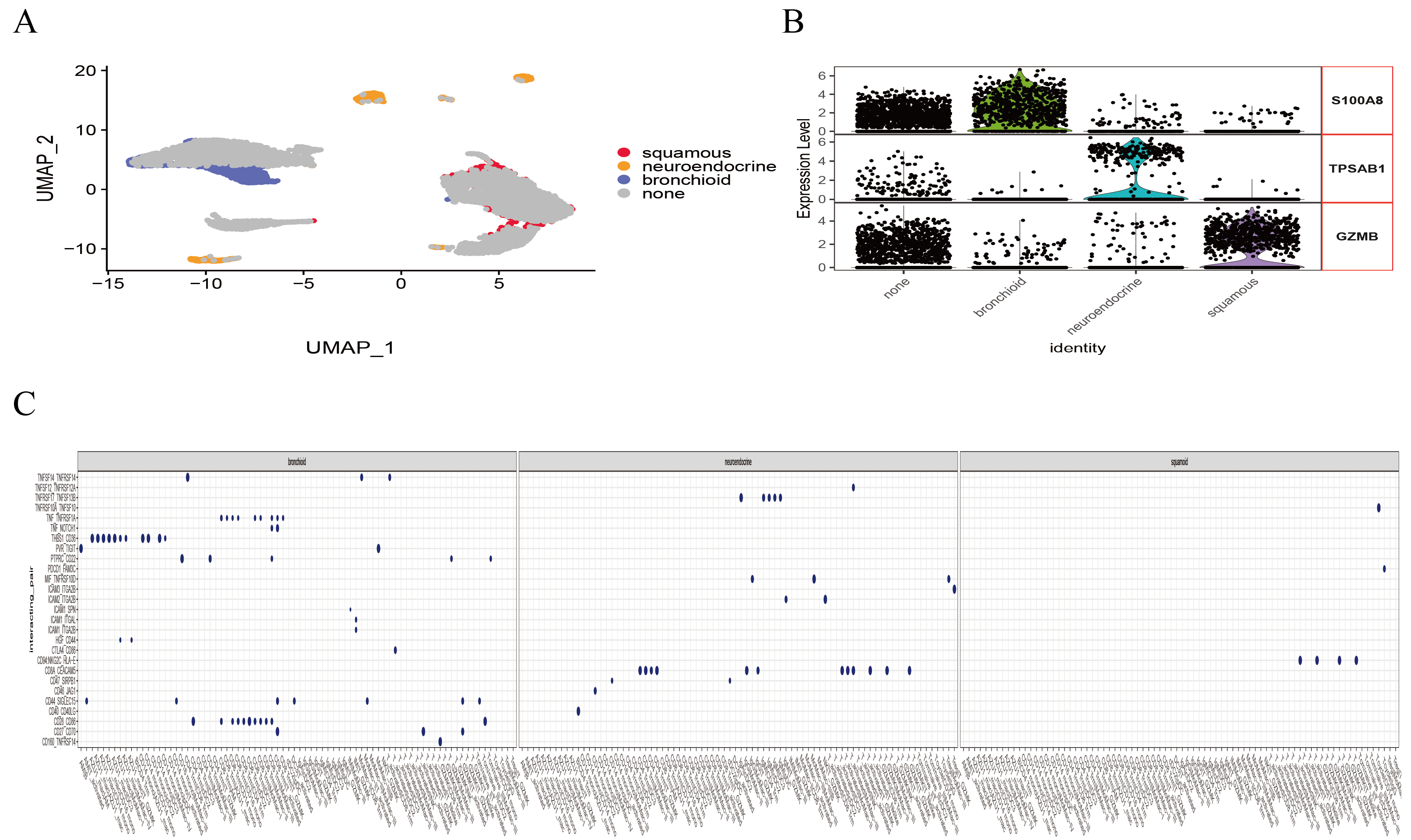

**Figure S3. Single-cell analysis among cluster-related cell subpopulation**

(A) Umap showing the relative distribution from bronchioid, neuroendocrine and squamoid predicted clusters. Note that “none” represents the low confidence or unrelated above clusters. (B) Boxplot plotting classical marker molecules among the above clusters. (C) Unique inter-cellular signals among the above clusters (The vertical axis represents ligand-receptor pairs, with larger dot representing lower p-value. Only incorporating mean expression greater than 1 and p-value less than 0.05).

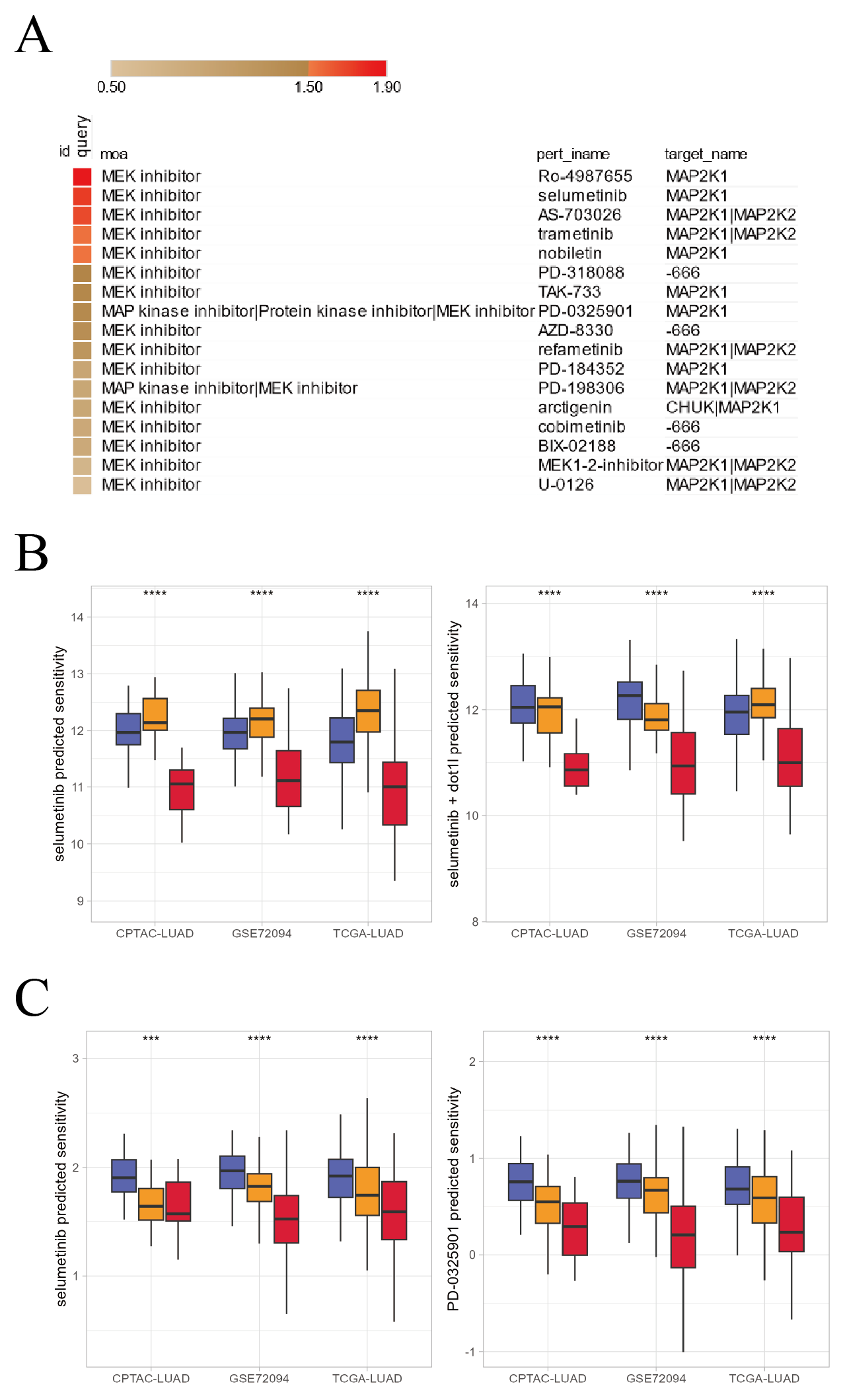

**Figure S5. MEK inhibitors sensitivity predicted by clusters**

(A) CMAP enrichment of MEK inhibitors using bronchioid-annotation cell lines. Enrichment greater than 1.5 was considered significant. (B) CCLE predicted sensitivity values in clusters using Kruskal-Wallis test. *, **, *** and **** representing p<0.05, p<0.01, p<0.001, respectively (left: selumetinib; right: selumetinib combination with DOT1L inhibitor). (C) GDSC predicted sensitivity values in clusters using Kruskal-Wallis test. *, **, *** and **** representing p<0.05, p<0.01, p<0.001, respectively (left: selumetinib; right: PD-0325901).

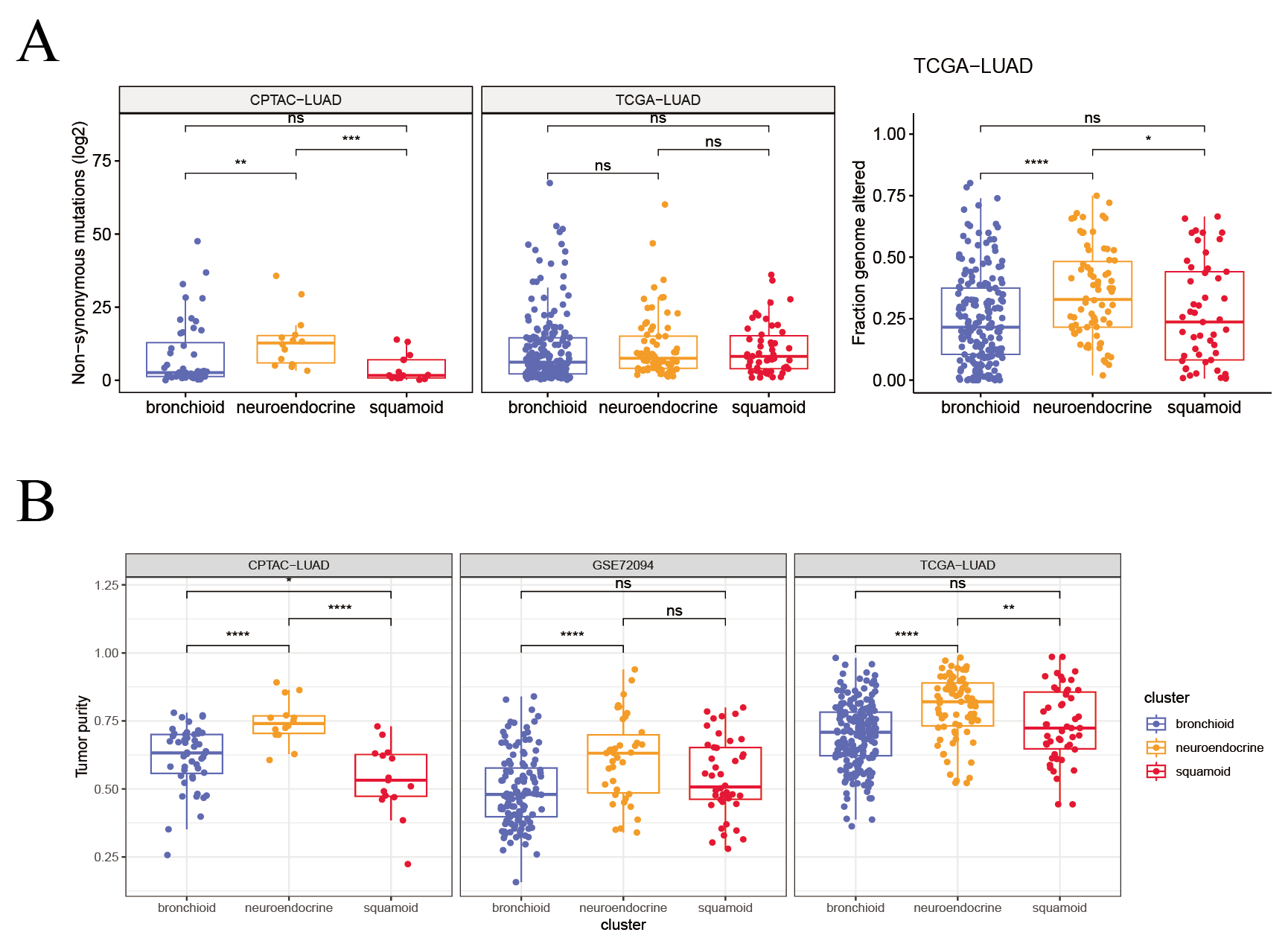

**Figure S6. Tumor burdens, fraction genome altered and tumor purity in clusters**

Boxplot showing (A) Non-synonymous mutations and fraction genome altered, and (B) tumor purity among three clusters in CPTAC-LUAD, GSE72094, and TCGA-LUAD cohorts using Kruskal-Wallis test. *, **, *** and **** representing p<0.05, p<0.01, p<0.001, respectively.

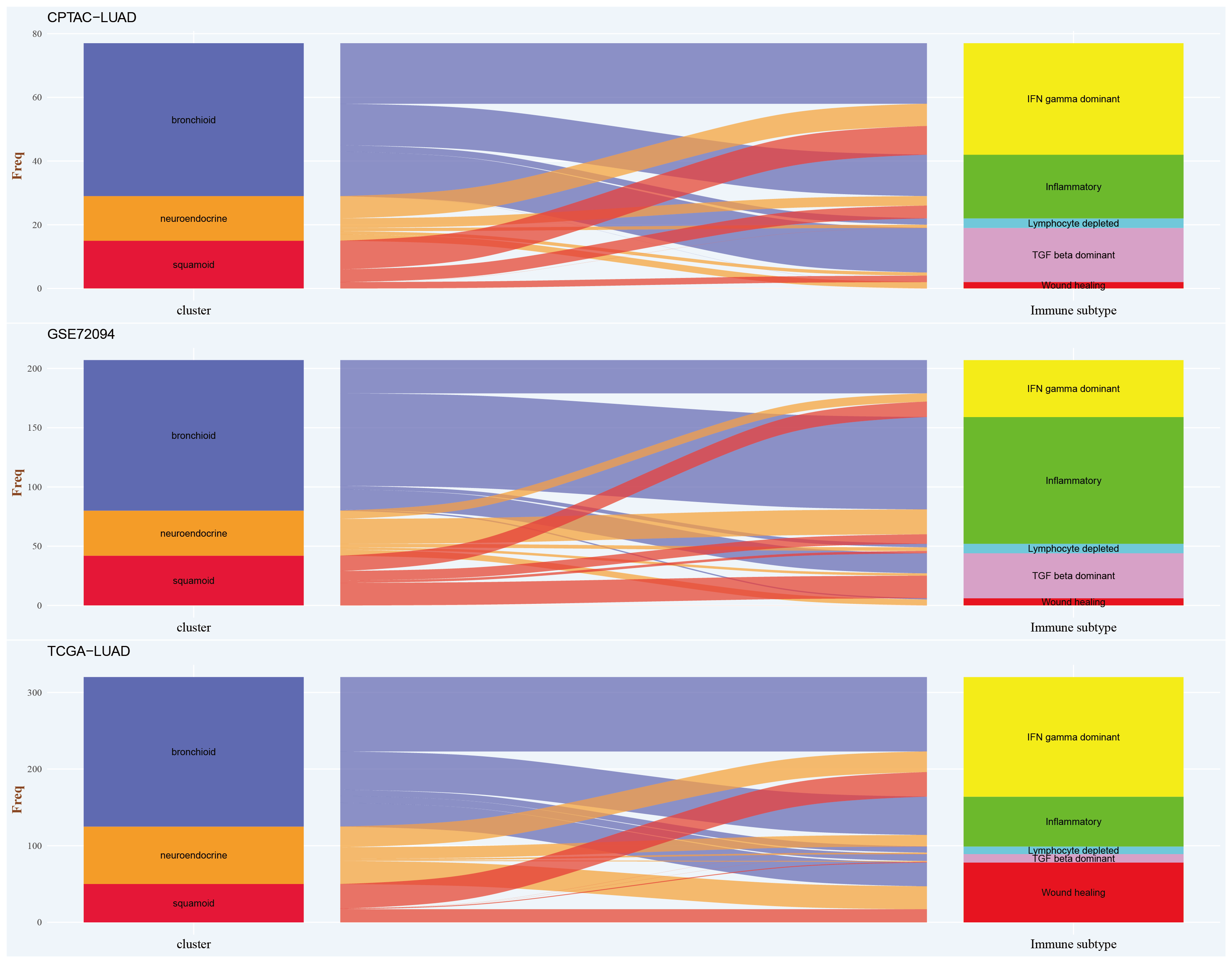

**Figure S7. Association between cluster and classical immunephenotype**

Sankey diagram displaying correspondence between cluster and immunophenotype in CPTAC-LUAD, GSE72094, and TCGA-LUAD cohorts (left: cluster; right: immunophenotype, the Y axes represent the number of tumor samples).

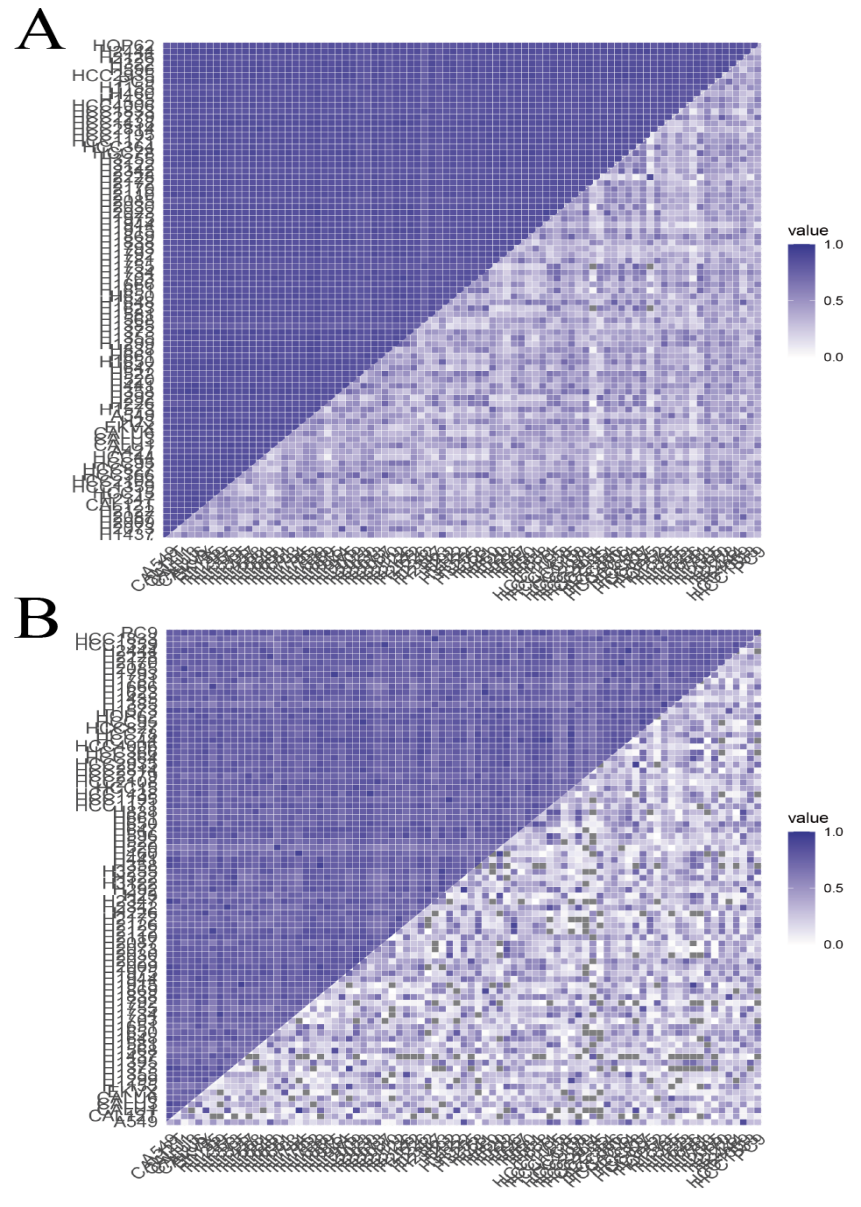

**Figure S8. Validation of cell lines correlations**

(A) Correlation matrix comparison using RNA expression (upper triangle: global gene; lower triangle: lineage-specific gene). (B) Correlation matrix comparison using RNA expression (upper triangle: immune pathway; lower triangle: 42-classifier).
